# Massive RNA editing in ascetosporean mitochondria

**DOI:** 10.1101/2023.07.06.548043

**Authors:** Akinori Yabuki, Chihaya Fujii, Euki Yazaki, Akihiro Tame, Keiko Mizuno, Yumiko Obayashi, Yoshitake Takao

## Abstract

Ascetosporeans are parasitic protists of invertebrates. As only two species of Mikrocytida, an ascetosporean subgroup, have ever been sequenced deeply and analyzed using cells isolated from infected organisms, it was shown that their mitochondria are functionally reduced and the organellar genome is lacking. However, molecular studies on other ascetosporeans have not been conducted, and whether reduced mitochondria is common in ascetosporeans remains unclear. In the present study, we established two cultures of Paradinida, another ascetosporean subgroup, and reconstructed their mitochondrial genomes. As they were compared with their RNA-seq data, massive A-to-I and C-to-U types of RNA editing were detected. Many editing sites are shared between two paradinids, but strain-unique sites also exist. As the mitochondrial genes are involved in the electron transfer system, their mitochondria are not functionally reduced, unlike that in Mikrocytida. Furthermore, we detected adenosine deaminase acting on RNA (ADAR), which is a key enzyme of A-to-I substitution, in paradinids as well as several other protists. Immunostaining showed that this ADAR is specifically localized in the mitochondria of paradinids, suggesting that A-to-I substitution in paradinid mitochondria is mediated by ADAR. These findings elucidated the functional diversity and evolutionary process of ascetosporean mitochondria as well as ADAR.

## Introduction

Ascetosporea (ascetosporeans) is a class of Endomyxa, Rhizaria, and all its species are parasites of invertebrates. Their cell cultures do not exist, and the life cycle of ascetosporeans is still unclear^1,2^. As some of their members cause serious damage to aquacultures, they are also important research targets in fishery sciences^3-5^. While five major groups are recognized in Ascetosporea, only two species of Mikrocytida have ever been sequenced deeply and analyzed using cells isolated from infected oysters and crabs^6, 7^. Pioneering studies showed that their mitochondria are functionally reduced to mitochondrion-related-organelles (MRO), and their organellar genomes are lacking. However, in-depth studies on other ascetosporeans have not been conducted yet, and whether MRO is common in ascetosporeans as a whole remains unclear.

RNA editing is an essential cellular function, resulting in RNA modification. Several types of RNA editing have been reported, and they play an important role in the alteration of functional proteins and non-coding RNA^8^. Adenosine-to-inosine (A-to-I) substitution is likely the most studied form of RNA editing. This type of RNA editing is mediated by adenosine deaminase acting on RNA (ADAR) in the metazoan nucleus, and ADAR mutations are associated with several diseases in humans, including prostate cancer and amyotrophic lateral sclerosis^9-11^. The absence of ADAR in the genomes of fungi and early branching opisthokonts suggests that ADAR evolved from adenosine deaminase acting on tRNA (ADAT), which is conserved in all eukaryotes derived from the metazoan ancestor, and then diversified into several subfamilies acquiring several functional motifs and domains in the metazoan evolution^12^. Although ADAR was exceptionally reported from *Symbiodinium* spp.^13^, its origin and evolution are poorly understood. A-to-I type of RNA editing has been reported from the mitochondria of several protists, such as diplonemids,^14, 15^ and dinoflagellates^16^; however, their mediating mechanisms were not understood.

In the present study, we established clonal cultures of Paradinida, which is another subgroup of Ascetosporea, and their mitochondrial genomes were sequenced. By analyzing them with their RNA-seq data, it was revealed that massive RNA editing occur in their mitochondria and they have ADAR targeting into mitochondria. As the mitochondrial function of Paradinida was predicted not to be reduced, there is diversity about mitochondrial structure and function in Ascetosporea. Further, mitochondrial targeting ADAR, which has never been expected, also illuminated the origin and functional diversity of RNA editing medicated by ADAR.

## Materials and Methods

### Sample acquisition and culturing

Initial water samples were collected from Tokyo and Suruga Bay (Table S1). A small aliquot of each sample was added to Hemi medium^17^ with a 5-μl/ml antibiotic cocktail (P4083, Merck, Darmstadt, Germany) and incubated under dark conditions at 19–20 °C. From the incubated samples, cultures of FC901 and SRM-001 were established by isolating a single cell using a glass micropipette, and the cultures were axenically maintained by inoculation in Hemi medium without antibiotics at 19–20 °C under dark conditions every two weeks. The absence of contaminating bacterial cells in the culture (i.e., axenic culture) was confirmed by careful microscopic observation and PCR using the extracted total DNA with universal bacterial primer set, i.e., 27f^18^ and 1492r^19^.

### Microscopy

Living cells of Paradinida spp. FC901 and SRM-001 were observed under a BX43 microscope (Olympus, Tokyo, Japan) equipped with a digital 4K camera, FLOYD-4K (Wraymer, Osaka, Japan). For scanning electron microscopy, the axenic cells that grew on a glass slide were fixed with 2.5% glutaraldehyde in the cultivation medium at 4 °C. The fixed cells were washed with 0.22-μm-filtered artificial seawater (FASW; 3.5% Rei-Sea Marine II; Iwaki Co. Ltd., Tokyo, Japan) and then postfixed with 2.0% osmium tetroxide dissolved in FASW for 2 h. The postfixed cells were dehydrated using a graded ethanol series, dried with a JCPD-5 critical point drying device (JEOL, Akishima, Japan), then coated with osmium using an OPC-80 osmium coater (Filgen, Nagoya, Japan). The specimens were imaged using a field-emission scanning electron microscope (Quanta 450 FEG; Thermo Fisher, Waltham, MA) operating at 5 kV.

### Sequencing analyses

Approximately 200 ml of mid-exponential phase cell cultures of Paradinida spp. FC901 and SRM-001 were centrifuged at 2,400 × *g* for 5 min. The cell pellets were frozen and sent to the sequencing company (Azenta, Tokyo, Japan), and the library reconstruction and sequencing analyses were conducted using default settings. The details of the analyses and the sequence outputs are summarized in Table S2.

The raw fastq data of DNA-seq were divided into 100 subsets using SeqKit^20^, and three subsets of each strain were subjected to the contig assembly using SPAdes 3.13^21^ with default settings. From each assembly data, a single possible mitochondrial genomic fragment was detected by BLASTN using the mitochondrial genome sequence of *Ophirina amphinema* (GenBank accession number: LC369600.1) as the query sequence. The detected sequences were identical among three subset analyses of each strain, while the starting position of each sequence differed. By comparing these sequences, a circular mitochondrial genome of FC901 and SRM-001 was reconstructed. The same assembly analyses were also conducted using the RNA-seq data. The obtained mitochondrial sequences, which were reconstructed from RNA-seq data, were subjected to annotation using MFannot (https://megasun.bch.umontreal.ca/apps/mfannot/) and compared with those assembled from DNA-seq data using Mesquite 3.10^22^.

For analyzing the transcriptome data, three RNA-seq datasets of Paradinida sp. FC901 were combined into a single dataset. The fastq data of each strain were subjected to contig assembly using SPAdes 3.13^21^ with the ‘--rna’ option. From the reconstructed contigs, their ADAR and ADAT sequences were searched by TBLASTN using the ADAR sequence of *Symbiodinium microadriaticum* (OLQ07757; E-value cut-off was set to 10^−10^). We also searched publicly available sequencing data (Table S3) for ADAR and ADAT sequences of other protists using the same approach. The detected sequences (e.g., ADAR and ADAT of *Phaeodactylum tricornutum*) were also used as the query in further searches for identifying more ADARs and ADATs. The obtained sequences were aligned with those of the metazoan ADARs and ADATs and then subjected to automated alignment using MAFFT v 7.471 with the ‘L-INS-’ option^23,24^. The aligned sequences were masked for the phylogenetic analysis using trimAl v1.4 with the ‘strict’ option^25^. This initial dataset contained all the detected sequences, including the partial short and highly divergent sequences, and only 94 positions were included in the phylogenetic analysis. The tree topology and branch lengths were inferred using the maximum likelihood (ML) methods using IQ-TREE 2.2.0^26^ with the LG+F+I+G4 model. The robustness of the ML phylogenetic tree was evaluated using a non-parametric ML bootstrap analysis with the LG+F+I+G4 model (100 replicates). We also conducted Bayesian phylogenetic analysis with the CAT + GTR model using PhyloBayes MPI v. 1.8a^27,28^. The analysis included two Markov chain Monte Carlo runs of 100,000 cycles with a ‘burn-in’ of 25,000 cycles. The consensus tree with branch lengths and Bayesian posterior probabilities were calculated from the remaining trees. Based on these findings, we revised the main dataset, excluding 14 partial and divergent ADAR sequences from an initial alignment. The main dataset was prepared using the same method used for the initial dataset and comprised 209 positions. The same methods were used to infer the phylogenetic tree and statistical support. Of the newly detected ADAR sequences, 12 sequences were retained in the main dataset and were subjected to motif identification by HMMER 3.3 (http://hmmer.org) along with the ADAR of Paradinida sp. FC901 against the Pfam database^29^.

The 18S rRNA gene sequences of Paradinida sp. FC901 and SRM-001 were determined using the DNA that was extracted with Qiagen DNeasy Plant Mini Kit (Qiagen, Hilden, Germany) from 20 ml of culture. The primers used were Euk1A^30^ and EukB^31^. The sequences were added to the alignment that was created based on the method proposed by Ward *et al*.^32^ and aligned using MAFFT v 7.471 with the default settings. The ML tree with the non-parametric bootstrap analyses of 1,000 replicates and the Bayesian tree were reconstructed using the same methods described earlier^33^.

### Western blot analysis

Approximately 200 ml of mid-exponential phase cell culture of Paradinida sp. FC901 was centrifuged at 2,400 × *g* for 5 min. The cell pellet was frozen and sent to Genostuff Co. Ltd. (Tokyo, Japan), where the following experiments were conducted.

The frozen cell pellet was homogenized in RIPA buffer (Fujifilm, Osaka, Japan) containing 1/100 (v/v) in a final volume of Protease Inhibitor Cocktail (Merck) for 30 min at 4 °C. After centrifugation at 18,000 × *g* for 5 min at 4 °C; the aqueous phase was recovered and utilized as the initial protein assay. The volume of the extracted proteins was measured using a BCA Protein Assay Kit (Thermo Scientific). The proteins (30 μg) were mixed with sample buffer (Thermo Scientific) and Sample Reducing Agent (Thermo Scientific) and then separated by 5% to 20% gradient polyacrylamide gel electrophoresis. The separated proteins were transferred onto a PVDF membrane (ATTO, Tokyo, Japan) and blocked for 1 h at room temperature with 0.1% TBST buffer containing 5% skimmed milk powder. The blotted membrane was incubated overnight at 4 °C with a polyclonal anti-ADAR antibody HPA051519 (Merck) diluted 1:500 with 0.1% TBST containing 5% BSA at final concentration. The membrane was washed four times in 0.1% TBST buffer and incubated for 1 h at room temperature with anti-rabbit IgG conjugated to HRP-linked antibody #7074 (Cell Signaling Technology, Danvers, MA) diluted 1:5,000 with 0.05% TBST containing 5% skimmed milk. After the membrane was rewashed four times in 0.05% TBST buffer, the bound antibodies were visualized using Immobilon (Merck) and recorded on C-DiGit (LI-COR, Lincoln, NE).

### Fluorescence assay

The cells of Paradinida spp. FC901 and SRM-001 were fixed with 4% paraformaldehyde in cultivation medium, centrifuged at 2,400 × *g* for 5 min, and then embedded in 1% agarose in FASW. The cells in the agarose gel were washed with FASW, dehydrated in a graded series of ethanol (30%, 50%, 70%, 90%, and 100%), and embedded in Technovit 8100 resin (Mitsui Chemicals, Tokyo, Japan) at 4 °C. Semi-thin sections (approximately 1-μm-thick) were cut using a glass knife mounted on an Ultracut S ultra-microtome (Danaher, Washington DC) and collected on a glass slide. The cell sections were treated with 2% block ace (KAC Ltd., Kyoto, Japan) in 1× PBS for 20 min at room temperature and then incubated with the anti-ADAR antibody HPA051519 (Merck) diluted 1:200 with PBS for 12 h at 37 °C. After incubation, the sections were incubated again with CF565-conjugated goat anti-rabbit IgG secondary antibody (1:200 dilution in PBS; Nacalai Tesque, Kyoto, Japan) for 2 h at room temperature and then stained with 1 μM Mito View Green solution (Biotium, Fremont, CA) and 4′,6-diamidino-2-phenylindole (DAPI) for 30 and 5 min, respectively. The sections were observed using a BX-51 light and fluorescence microscope (Olympus) with UV (Ex, 330–385 nm; Em, > 400 nm), FITC (Ex, 470–495 nm; Em, 510–550 nm), and CY3 (Ex, 530–570 nm; Em, 573–648 nm) filter sets for DAPI, antibodies, and Mito View Green, respectively.

The living cells of Paradinida sp. FC901, as a representative paradinid, were stained with 1 μM Mito View Green solution (Biotium) and DAPI on a glass slide for 30 min and 5 min, respectively, and observed using a BX-51 light and fluorescence microscope with UV and FITC filter sets.

## Results and Discussion

### Paradinid culture, mitochondrial genome, and RNA editing

Two ascetosporean amoebae, namely, Paradinida sp. FC901 and Paradinida sp. SRM-001 (Fig. 1, Fig. S1, S2), were established as clonal and axenic cultures. They phylogenetically belong to Paradinida (paradinids), Ascetosporea, Rhizaria, in the 18S rRNA gene tree (Fig. 1). All species belonging to Ascetosporea are known to be parasitic organisms; thus, Paradinida sp. FC901 and SRM-001 may also exhibit parasitism in natural environments, as do other ascetosporean species.

**Figure 1.**
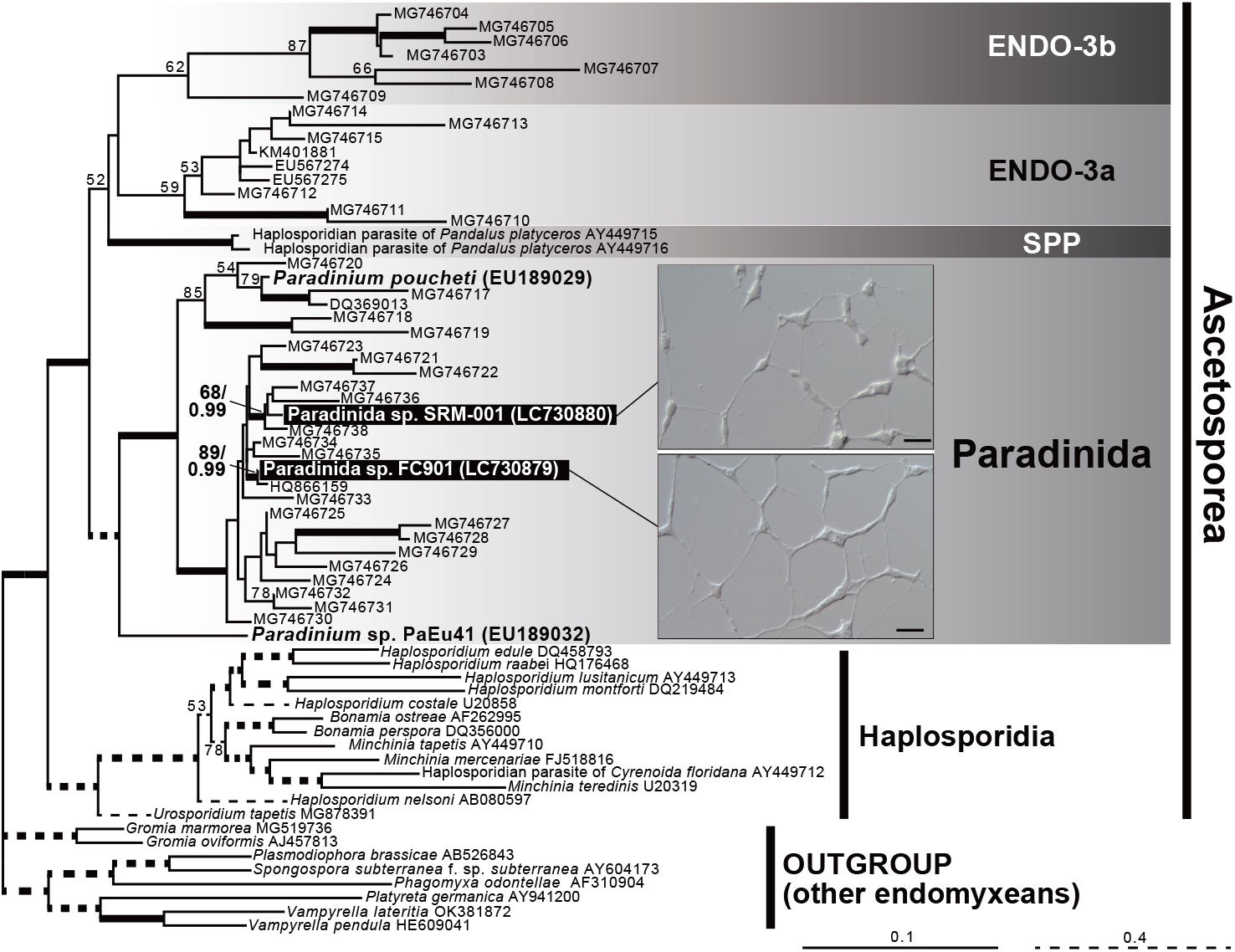
Phylogenetic tree of endomyxean 18S rRNA gene sequences with light microscopic images of Paradinida spp. FC901 and SRM-001. Only maximum likelihood (ML) bootstrap values > 50% are shown. The branches supported by > 0.95 of Bayesian posterior probability are shown in bold lines.

However, any specific details, including their host organism(s), remain unknown. As Paradinida spp. FC901 and SRM-001 are the first ascetosporean cultures, and they are maintained axenically; substantial progress has been achieved in the process of obtaining data from ascetosporean parasites.

By analyzing DNA-sequencing (seq) data, circular mitochondrial genomes were successfully reconstructed, and their lengths were 23,048 and 20,099 bp in FC901 and SRM-001, respectively (Fig. 2A). However, the coding regions in their genomes are finely fragmented by many unexpected stop codons, suggesting that they are either pseudogenes or involved in RNA editing. Hence, to confirm these possibilities, RNA-seq on both paradinids was conducted, and the obtained data were compared with DNA-seq data. The results of sequence comparison showed many adenosines and cytidines were switched to guanosines and uridines in RNA-seq data, respectively (Fig. 2B). As A-to-I substitution is the most common type of RNA editing and inosine is recognized as guanosine in reverse transcription, we considered that the paradinids also possess A-to-I substitutions in addition to C-to-U substitution in their mitochondria. Interestingly, many substitution sites are shared between FC901 and SRM-001, indicating that many of the editing sites existed previously in their common ancestor. Nonetheless, several strain-unique editing sites also existed (Fig. 2B), indicating that the acquisition of additional editing sites still progresses or progressed until just recently. All genes except *trnH* and *orf179* have editing sites, while the rate of editing varies among the genes (Fig. 2B). A-to-I substitution is more abundant than C-to-U substitution in all genes and the gene with the highest rate is *atp9* of SRM-001 (7.89%). The rate in all coding regions is 2.08% in FC901 and 2.37% in SRM-001 (see Supplementary Information; Table S4), which are values comparable to those of other protists possessing organellar RNA editing^34-37^, but much lower than the highest rate (12.7%) of a diplonemid *Namystynia karyoxenos*^38^. As any conserved and/or pattern sequences around the editing sites could not be detected, it is entirely unclear how those sites are specifically recognized for editing.

**Figure 2.**
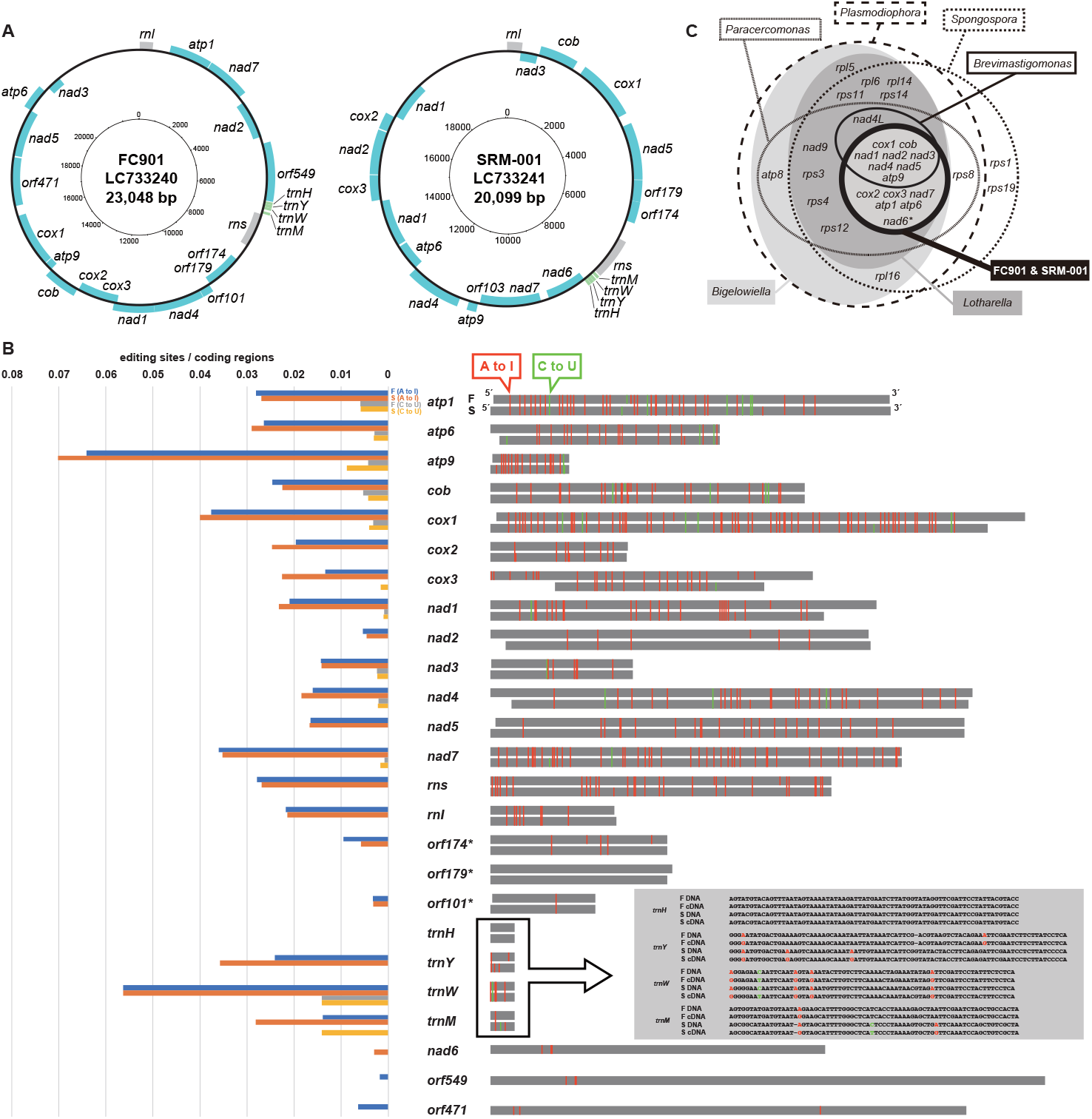
Summary of mitochondrial genome of FC901 and SRM-001 **A**. The mitochondrial genome map of Paradinida spp. FC901 and SRM-001. Protein-coding regions are shown in pale blue. Transfer RNA genes and other structural RNA genes (i.e., *rns* and *rnl*) are shown in green and gray, respectively. **B**. Summary of the RNA editing found in the mitochondria of FC901 and SRM-001. The left bars indicate the editing rate of each gene detected on the mitochondrial genome of FC901 and SRM-001. Blue, orange, gray, and yellow bars indicate the rate of A-to-I substitution in FC901 and SRM001 and the rate of C-to-U substitution in FC901 and SRM-001, respectively. Right-side bars indicate the length of each gene, and the editing sites are indicated by red (A-to-I substitution) or green (C-to-U substitution) vertical lines. The editing sites shared by FC901 and SRM-001 are indicated by the single connecting lines. **C**. Venn diagram summarizing the protein-coding genes on the mitochondrial genomes of related rhizarians. Only *nad6* is absent on the mitochondrial genome of FC901 but exists on that of SRM-001.

The gene repertory in the mitochondrial genome of the two paradinids is very similar, but *nad6* was only detected from SRM-001 (Fig. 2A, C). As only four species of tRNA (i.e., *trnH, trnY, trnW*, and *trnM*) are encoded on their mitochondrial genomes, we considered that other tRNA species were encoded on their nuclear genomes and transported into the mitochondria, as reported in other protists^e.g., 39-42^. The number of protein-coding genes is 13 in FC901 and 14 in SRM-001, and all of them have been reported from the mitochondrial genomes of other rhizarians sequenced previously (Fig. 2C); these genes are all functionally involved in the electron transport chain. Notably, because the genes coding for proteins seen in complex I, III, IV, and V were detected, we believe that the mitochondria of paradinids have a normal function to synthesize ATPs, and they are not functionally reduced to MROs, which is not consistent with that of Mikrocytida^6,7^. The function of ascetosporean mitochondria, including MRO, may be different, lineage by lineage, by the level of adaptation to the parasitic lifestyle.

### Detection of ADAR in paradinids and possible origin in LECA

A-to-I and C-to-U substitution types of RNA editing are found in not only protein-coding genes but also structural genes of paradinids. These types of RNA editing are also reported from diplonemids^38^, but the editing in the transfer RNAs is unique in paradinids. Since paradinids and diplonemids are phylogenetically distant from each other, their RNA editing was likely established independently. The mechanisms involved in RNA editing of diplonemids have not yet been elucidated, and the roles of key enzymes, i.e., ADAR for A-to-I substitution and APOBEC for C-to-U substitution, are not understood. By contrast, we successfully detected ADAR sequences from paradinids using TBLASTX (E-value cut-off was set to 10^−10^) with a *Symbiodinium* ADAR sequence (OLQ07757) as the query. APOBEC was not found in the paradinids investigated in this study.

Furthermore, as we searched for ADAR sequences from the publicly available sequencing data (Table S3), 28 possible ADAR sequences from 24 protist species in total were detected. Since 14 out of the 28 sequences were partial and/or highly divergent, their assignment as ADAR is not yet conclusive. By contrast, the other 14 sequences from 13 protists, including two paradinids, form a clade with moderate statistical support, while the sister relationship between protistan ADARs and the metazoan ADARs is not well supported (Fig. 3). In addition to this phylogeny, ADAT can be found in 7 of 11 protist species that possess ADAR (Table S3). Although the double-strand RNA binding motif, a key motif of metazoan ADARs, is absent from protistan ADARs (Fig. S3), they can be distinguished from ADAT, and it is reasonable to consider that protistan ADARs belong to the ADAR family. As the protists possessing ADAR belong to phylogenetically divergent lineages, ADAR likely evolved much earlier in the history of eukaryotes than previously thought and may have originated in the LECA. The secondary loss of ADAR possibly occurred independently at the base of fungi, as fungi do not have ADAR^43^.

**Figure 3.**
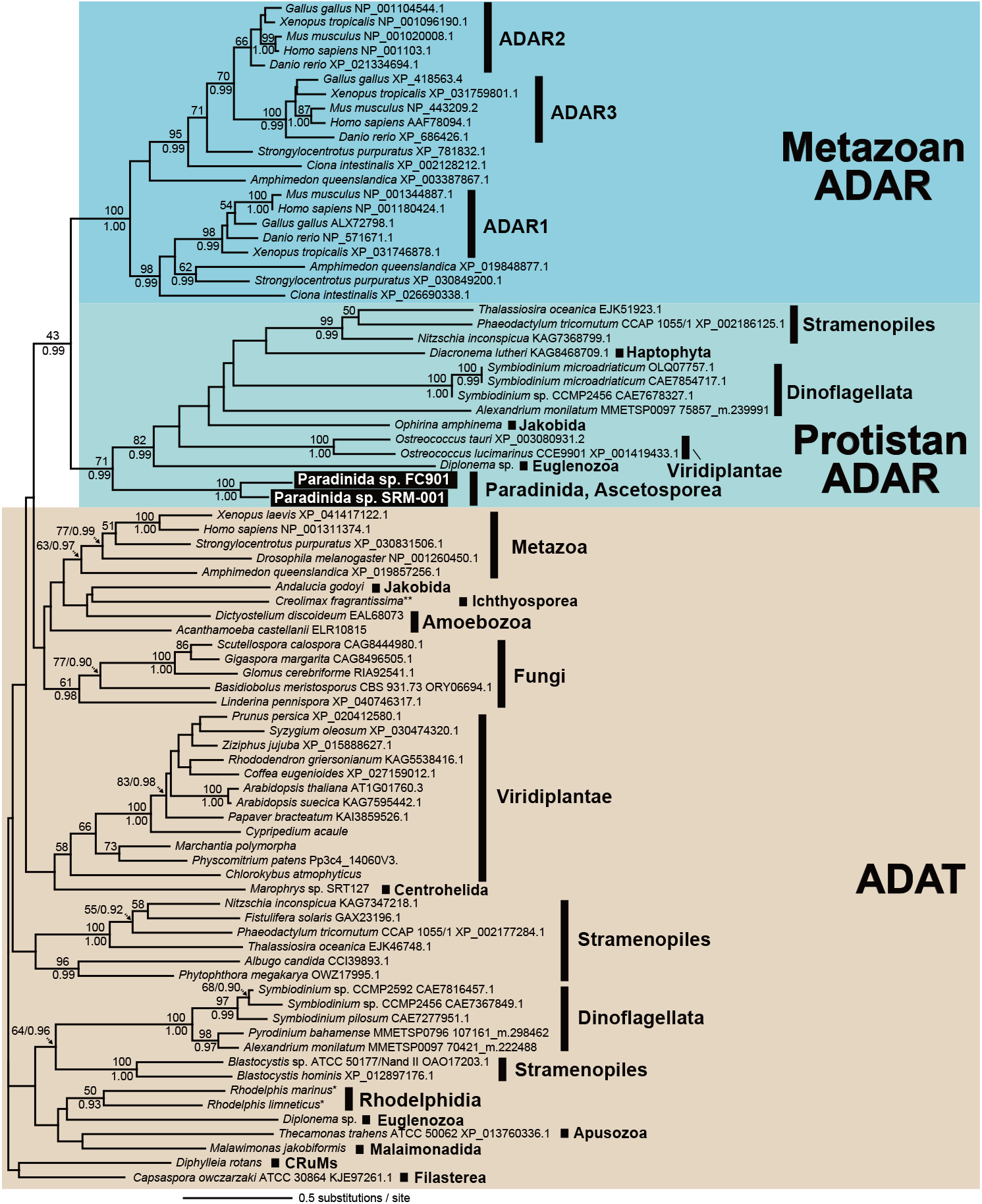
Phylogenetic tree of eukaryotic adenosine deaminase acting on RNA (ADAR) rooted with eukaryotic adenosine deaminase acting on tRNA (ADAT). The maximum likelihood (ML) trees inferred from ADAR–ADAT alignments are shown. ADAT sequences (and branches) are brown. Metazoan ADAR is blue, and protistan ADAR is light green. Only ML bootstrap values/Bayesian posterior probabilities equal to or > 50%/0.90 are shown.

### Subcellular localization of paradinid ADAR and its role in mediating mitochondrial RNA editing

It may be reasonable to consider that paradinid ADAR is involved in mitochondrial RNA editing; however, the specific ADAR in mitochondrion has not been reported to date. All of the ADAR for which subcellular localization was studied belong to metazoans, and they are all localized in the nucleus except for a single isoform found in the cytosol. The structures stained by anti-human-ADAR antibody HPA051519 clearly overlap with the mitochondria stained by MitoView (Fig. 4A-F) in two paradinids; western blotting analysis was used to study its specificity for the extracted proteins of FC901 (Fig. S4). The structures stained by MitoView are also consistent with the distribution pattern of the mitochondria stained by MitoTracker (Fig. S5). These findings are very consistent with the fact that A-to-I substitutions occur in the mitochondria of paradinids; hence, we consider these ADARs probably contribute to RNA editing in their mitochondria.

**Figure 4.**
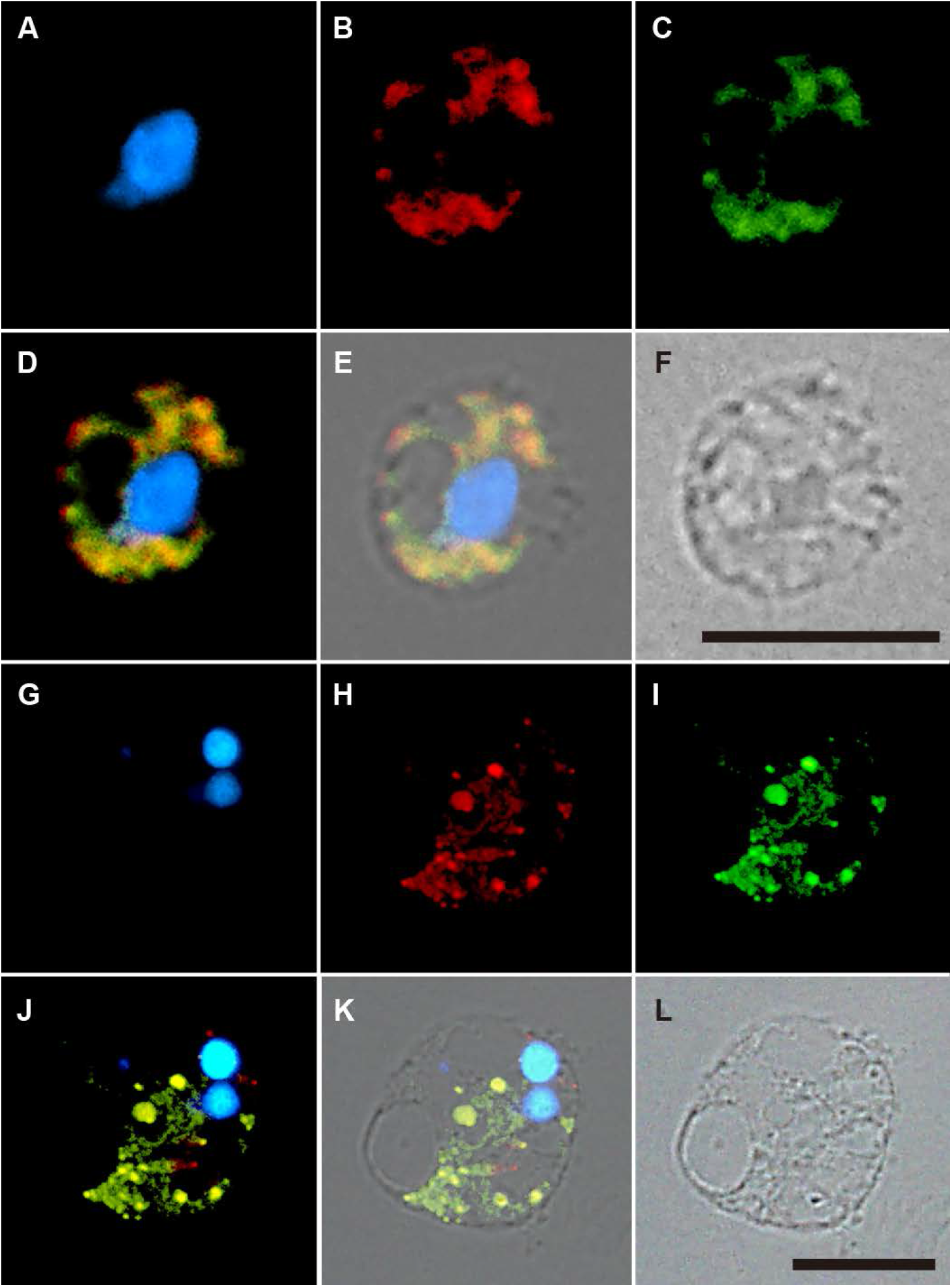
Subcellular localization of paradinid’s ADAR. **A-F**: FC901, **G-L**: SRM-001. Cell sections are stained by DAPI (**A, G**), anti-ADAR antibody (**B, H**) and Mito View (**C, I**). Differential interference contrast image of cell sections (**F, L). D**. Merged view of Fig. 4A–C. **E**. Merged view of Fig. 4A–C and F. **J**. Merged view of Fig. 4G–I. **K**. Merged view of Fig. 4G–I and L. Bars = 10 μm.

The subcellular localization and function of other protistan ADARs remain unclear. Of the protists possessing ADAR, only dinoflagellates, diplonemids, and paradinids possess A-to-I substitution in their mitochondrial genes^14-16^. Although the mitochondrial genome of the other protists that was shown to have ADAR in this study has already been reported^44-46^, the possible existence of A-to-I substitution in their mitochondria was neither detected nor suggested. Their ADARs might contribute to the editing of non-coding RNA in their mitochondria; however, it may be more reasonable to consider that their ADARs function in the nucleus, as do the metazoan ADARs. Ancestral paradinids’ ADAR may also have been localized in the nucleus and then underwent changes to function in the mitochondrion. As we could not detect ADAR from the nucleus of paradinids, the original function of ADAR in paradinids’ nucleus may have not been crucial.

### Collateralization of life for diversification (COLD) hypothesis explains the advantage of complex mitochondrial RNA editing

The RNA editing in paradinids’ mitochondria is very complex with respect to the number of editing sites. More than 2% of the total coding region is involved in the editing process. If the mitochondrial genomes only encode proteins that do not require RNA editing, the underlying processes and molecular machineries for RNA editing are unnecessary. Paradinids rather seem to have survival constraints in possessing RNA editing; paradinids must keep their organellar editing function to have operational proteins in their mitochondria. This complex characteristic from which adaptive advantages cannot be explained may have been established due to constructive neutral evolution (CNE). CNE posits that complexity can increase for a long period, even if the complexity itself is neutral or slightly negative for the survival of each organism^47, 48^. However, here, we found and proposed a possible indirect advantage to possess organellar RNA editing.

While organellar RNA editing may be just extra steps for usual gene expression, the protists possessing it (e.g., diplonemids and paradinids) are distributed with high lineage diversity in oceans 32,49. In other words, they succeeded in diversifying, and we can, thus, hypothesize that organellar RNA editing may have partially contributed to their diversification. Mutations in any kind of genome always occur with a certain probability. If a lethal mutation occurs in the mitochondrial genome, the individual carrying such a mutation cannot survive. However, if the organism has RNA editing activity in the mitochondria and can mitigate the effects of such lethal mutations by RNA editing, then such mutations may not be lethal. In the case of paradinids, some mutations, which are lethal and involved in the substitution of adenosines or cytidines from other nucleotides, can be masked by RNA editing activity. Although that organism must keep the RNA editing function for survival, the lethal mutation in the mitochondrial genome is less of a constraint factor for the survival and diversification of that organism and its descendants. However, as the constraint on the mutation gets relaxed in the organismal lineage, the complexity (i.e., the number of editing sites) continues to increase, and it is impossible to return to the original non-complex state. Until the day when the complexity reaches a limitation, RNA editing can mask some of the lethal mutations in the mitochondrial genome and help organismal diversification. However, when the complexity reaches a plateau and the RNA molecules cannot be modified by the existing mechanisms, the organisms cannot survive any longer. In other words, the organisms possessing the organellar RNA editing may have some advantage for diversification instead of stocking the potential risk for future extinction (Fig. 5). Here, we propose this evolutionary scenario as the COLD hypothesis. The COLD hypothesis is based on CNE but differs from CNE as an adaptive indirect advantage is found in a limited time. Although the corrective function of RNA editing was also indicated in the previous studies ^e.g., 37^, the link between its function and organismal diversification has not been discussed. In the long history of this planet, the diversification and extinction of various organisms have occurred continuously, and some of these events might be explained by the COLD hypothesis.

**Figure 5.**
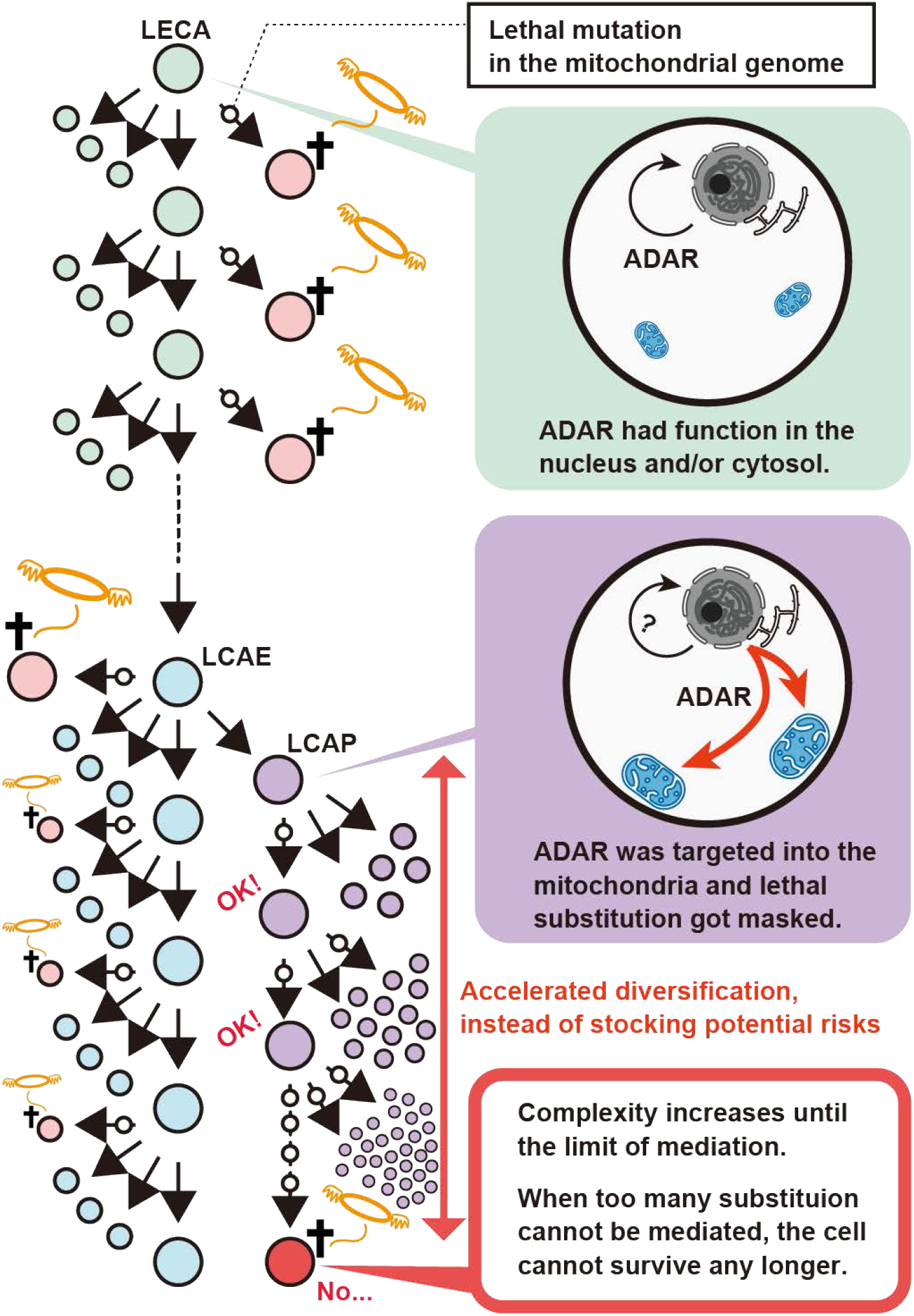
Schematic summary of the COLD hypothesis. LECA, last eukaryotic common ancestor; LCAE last common ancestor of endomyxeans; LCAP, last common ancestor of paradinids. White circles indicate the occurrence of a lethal mutation in the mitochondrial genome.

## Data availability

Mitochondrial genomes for Paradinida spp. FC901 and SRM-001 are available under GenBank accession numbers LC733240 and LC733241, respectively. The raw sequencing data for genome reconstruction and confirmation of RNA editing is available under GenBank BioProject accession number PRJDB14367. Their 18S rRNA gene sequences are available under GenBank accession numbers LC730879 (FC901) and LC730880 (SRM-001). The ADAR sequences that were newly detected and analyzed in this study as well as the datasets, can be found in online repositories, Dryad https://doi.org/10.5061/dryad.mcvdnck4z.

## Acknowledgments

We thank the Captain and crew of R/V Hokuto (Tokai University) and all the members of the SURUME (Suruga Bay Research for Understanding Marine Ecosystem) Project for their kind support on sampling.

